# Nerpa 2: probabilistic linking of biosynthetic gene clusters to nonribosomal peptides

**DOI:** 10.1101/2024.11.19.624380

**Authors:** Ilia Olkhovskii, Aleksandra Kushnareva, Azat Tagirdzhanov, Alexey Gurevich

## Abstract

**Motivation:** Nonribosomal peptides (NRPs) are bioactive microbial metabolites with high pharmaceutical potential. Although genome mining enables large-scale detection of biosynthetic gene clusters (BGCs) predicted to encode NRPs, reliably linking these clusters to their chemical products remains challenging due to the flexible and heterogeneous organization of NRP assembly pathways.

**Results:** We present Nerpa 2, a probabilistic framework for accurate and scalable linking of NRP BGCs to candidate chemical structures. The method represents assembly lines as hidden Markov models (HMMs) that capture uncertainty and alternative biosynthetic routes. On curated datasets of experimentally validated BGC–product pairs, our tool outperforms existing methods in linking accuracy and pathway reconstruction. When applied to large genome mining datasets, Nerpa 2 efficiently identifies BGCs likely associated with known compounds and highlights potential producers of novel chemistry.

**Availability and implementation:** Nerpa 2 is freely available at https://github.com/gurevichlab/nerpa.

## 1. Introduction

Nonribosomal peptides (NRPs) are bioactive microbial natural products that include clinically important antibiotics and other therapeutics (Agrawal *et al*., 2017; Süssmuth and Mainz, 2017). Advances in genome sequencing have enabled large-scale detection of biosynthetic gene clusters (BGCs) encoding NRPs, but linking these clusters to the chemical structures they produce remains a central challenge in genome mining (Medema *et al*., 2021).

The difficulty arises from the modular architecture of NRP synthetases: while each module typically incorporates one amino acid, substrate-selecting adenylation (A) domains can be promiscuous, module activation may deviate from gene order (non-collinearity), and modules can be skipped or reused (Süssmuth and Mainz, 2017; Juguet *et al*., 2009). In addition, downstream enzymatic modifications further diversify the final product (Sieber and Marahiel, 2003). These features complicate direct sequence-to-structure matching.

Here we present Nerpa 2, a complete methodological rewrite of our previous tool (Kunyavskaya *et al*., 2021). The new version replaces a dynamic programming–based alignment scheme with a probabilistic hidden Markov model (HMM) framework decoded using the Viterbi algorithm. This formulation explicitly models module skipping, insertions, and substrate uncertainty. On curated benchmarks, Nerpa 2 improves top-10 BGC–NRP linking accuracy by roughly one third and reduces alignment errors by more than fourfold compared to existing methods. The tool also scales to hundreds of millions of BGC–NRP comparisons, supporting large-scale discovery efforts.

## 2. Materials and methods

### 2.1. Pipeline overview

Nerpa 2 links BGCs to candidate NRP structures by combining genome mining, chemical structure decomposition, and a probabilistic alignment framework (Fig. 1).

**Figure 1:**
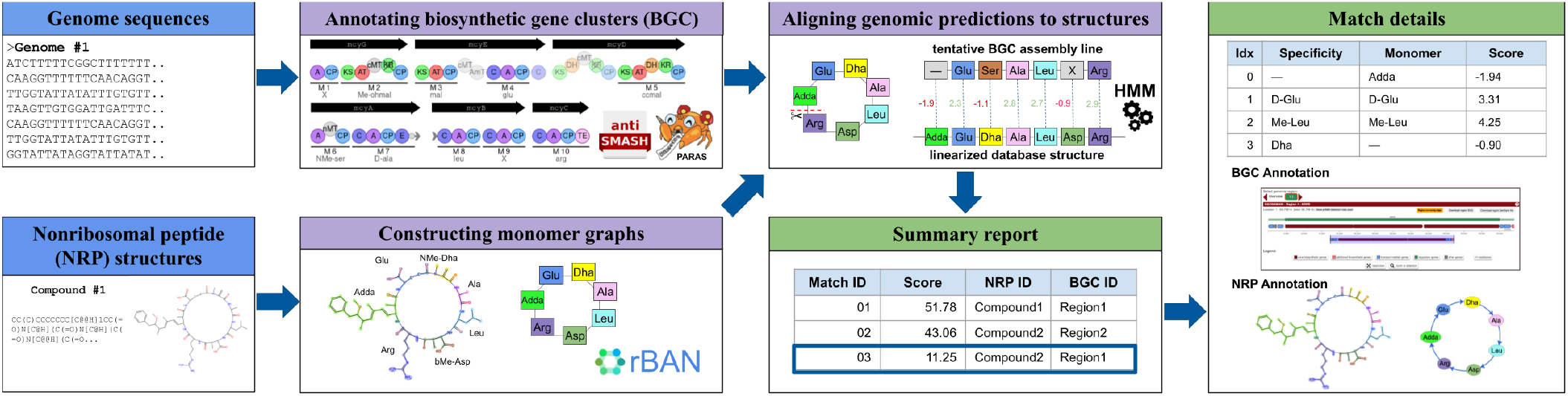
Nerpa 2 pipeline. NRP-related BGCs are identified from genome sequences, and A-domain specificities are predicted. NRP chemical structures are decomposed into monomer graphs and linearized. The tool represents candidate biosynthetic assembly lines as HMMs and aligns linearized NRP sequences against them. Top-scoring BGC–NRP matches are reported in machine-readable and interactive outputs.

Genome sequences are analyzed with antiSMASH (Blin *et al*., 2025) to detect NRP-related BGCs and annotate their modular organization. A-domain substrate specificities are predicted using PARAS (Terlouw *et al*., 2025) and converted into probability distributions over supported NRP building blocks (hereafter referred to as *monomers*). Based on these predictions and genomic context, candidate biosynthetic assembly lines are inferred and represented as HMMs.

In turn, NRP chemical structures are decomposed into monomer graphs using rBAN (Ricart *et al*., 2019) and transformed into candidate linear monomer sequences. Each sequence is evaluated against each BGC-derived HMM using a log-odds scoring scheme relative to a null model, and top-scoring BGC–NRP matches are retained.

### 2.2. Representation of BGCs and NRPs

#### 2.2.1. Monomer set

Nerpa represents both BGCs and NRPs using a common set of monomers. Each monomer is described as a triplet (*core residue, methylation, stereochemistry*), where the *core residue* corresponds to the amino acid selected by the associated A domain and the remaining two fields capture optional post-selection modifications. *Methylation* indicates the presence of a methyl group (e.g., `NMe`, `OMe`), and *stereochemistry* indicates *L* or *D* configuration.

We currently support 30 core residues derived from the common substrates modeled by PARAS, together with an additional “`unknown`” placeholder. Building blocks identified by rBAN in NRP structures are classified as A-domain–related or A-domain–independent (e.g., polyketide fragments). The former are mapped to supported core residues using a curated conversion table, whereas the latter are removed during structure preprocessing.

#### 2.2.2. BGC module emission modeling

For each A domain detected by antiSMASH, Nerpa 2 applies the PARAS substrate specificity predictor trained on 34 common substrates (Terlouw *et al*., 2025). PARAS outputs a probability distribution over substrates, which are mapped to the corresponding core residues in the monomer set; probabilities of substrates mapping to the same residue are aggregated.

Because PARAS confidence scores do not directly correspond to incorporation probabilities, Nerpa 2 performs empirical calibration using curated BGC–NRP alignments (Section 2.5). Calibrated values are normalized to obtain a valid probability distribution over core residues (Supplementary Note S1).

Experimentally validated A-domain 8Å signatures (Röttig *et al*., 2011) are incorporated via a lookup dictionary. If a detected A domain matches a known signature, the corresponding core residues are assigned probability 1 (and all others 0) prior to normalization. This allows validated specificities, including promiscuous cases with multiple supported residues, to be incorporated directly into the model.

Each BGC module is thus represented as a probability distribution over monomers, assuming independent selection of core residue, methylation status, and stereochemistry. Probabilities of the post-selection modifications are determined by the presence of methylation and epimerization domains within the module.

#### 2.2.3. NRP linearization

NRP structures are represented by rBAN as monomer graphs, with vertices corresponding to monomers and edges to chemical bonds. These graphs are simplified by removing non-peptide bonds and discarding vertices that are not part of the peptide backbone.

For components that are simple cycles, Nerpa 2 generates a number of linearizations by breaking the cycle at each possible position. All other components are linearized by identifying a path that visits each monomer once (Hamiltonian path). For multi-component structures, Nerpa 2 tries all permutations for all choices of linearizations per component.

The resulting linearizations define candidate monomer sequences used for subsequent probabilistic alignment.

### 2.3. Biosynthetic assembly-line modeling

#### 2.3.1. Construction of assembly lines

As module activation order can vary, Nerpa 2 generates a number of candidate assembly lines for each BGC. Nerpa 2 takes into account domain composition (e.g., starter condensation and termination thioesterase domains indicate probable start and termination points), as well as changes in the strand orientation. (Supplementary Note S2).

Each candidate assembly line is represented as a sequence of A-domain-containing modules, each represented by the specificity distribution of core residues, modification annotations, and relevant genomic context.

#### 2.3.2. HMM formulation

For each candidate assembly line, Nerpa 2 constructs an HMM that models possible biosynthetic paths consistent with the inferred module activation order (Fig. 2). The model contains explicit INITIAL and FINAL states and is organized as a chain of module-specific subgraphs, one per A-domain-containing module.

**Figure 2:**
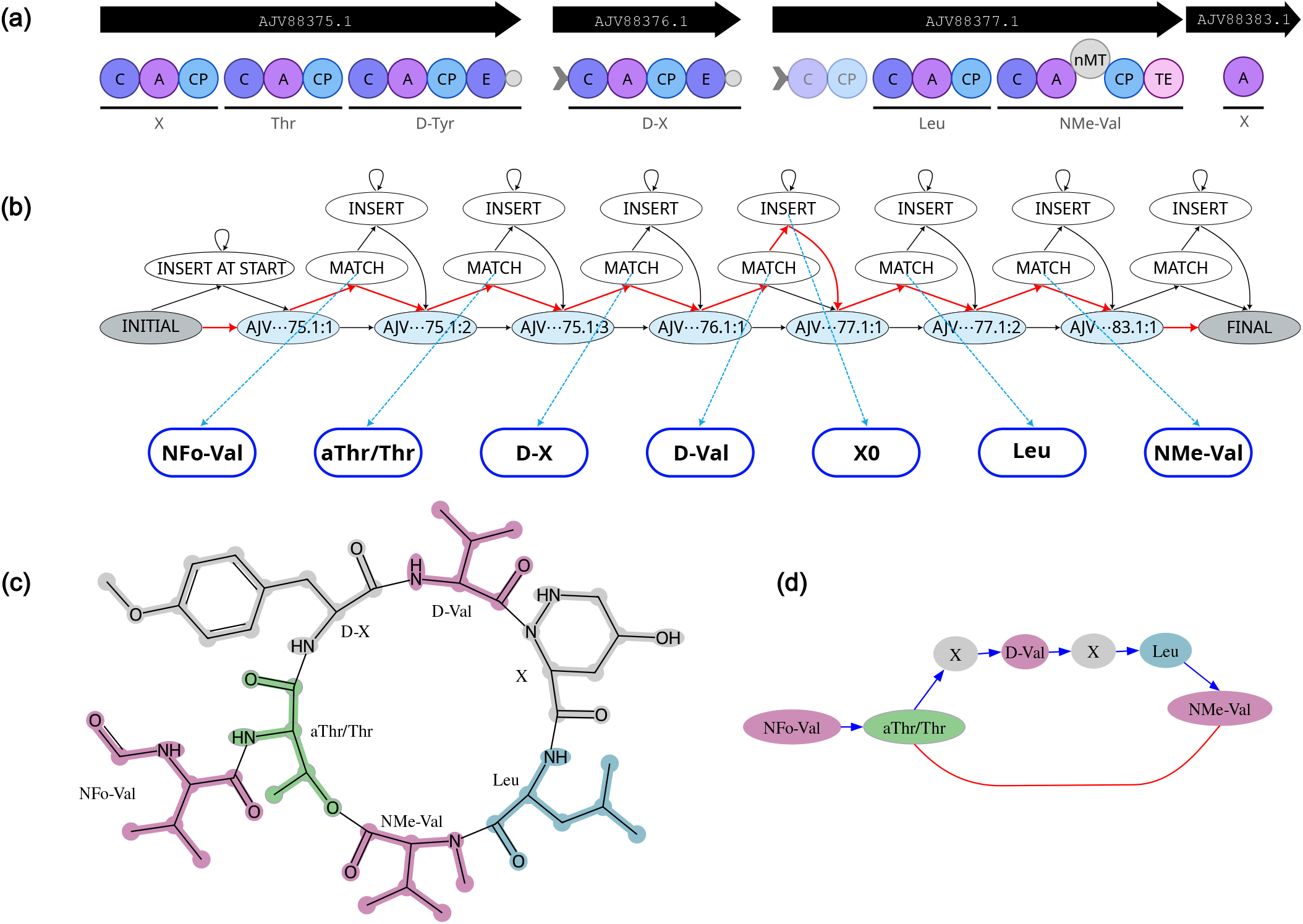
HMM formulation and example alignment. (a) Annotated BGC with genes (black arrows) and domains (C, A, CP, E, TE). The most likely A-domain specificity with modifications is indicated below the horizontal lines defining module borders; X denotes uncertain predictions. (b) Corresponding HMM (simplified), where each A-domain-containing module is represented by a subgraph centered on blue silent routing states. (c) Example NRP chemical structure annotated by rBAN and (d) its monomer graph representation. Blue (red) edges denote peptide (non-peptide) bonds; X labels are used for monomers not recognized by rBAN. In (b), blue dashed edges indicate emissions, and the most probable state path for the observed NRP monomer sequence (Viterbi decoding) is highlighted in red. The example is based on BGC0001214 from MIBiG and the corresponding marformycin E structure (Liu *et al*., 2015).

Within each subgraph, MATCH state emits monomers according to the module-specific emission distributions, while INSERT state emits monomers according to the background frequencies to capture incorporation events not explained by the core assembly-line modules. All the remaining states are silent.

The canonical path corresponds to sequential module activation, whereas alternative transitions enable non-collinear behaviors such as module skipping and insertion events.

Transition probabilities depend on module-level genomic context. For example, insertion transitions are favored when polyketide domains are located adjacent to a module, reflecting potential polyketide–NRP hybrid architecture, whereas skip transitions are more likely for modules consisting of only a single A domain, which are less likely to function as complete biosynthetic modules.

### 2.4. Alignment and scoring

To assess the likelihood of a BGC-NRP link, Nerpa 2 scores each linearization of the NRP against each HMM derived from the BGC, and against the null model.

The null model is independent of any BGC and assumes that each NRP monomer was generated from the same background monomer distribution. The latter was derived from the monomer frequencies in Norine (accessed on 17.01.2025), a curated database of known NRP structures (Flissi *et al*., 2020). Core-residue and modification probabilities (methylation and epimerization) were estimated independently, and the probability of each monomer was approximated as the product of the probabilities of its components.

The likelihood of an NRP linearization under the null model is thus the product of the background probabilities of its monomers.

The score of a match between an HMM and an NRP linearization pair is defined as the log-odds ratio between the probability of the most probable state path in the HMM, inferred using the Viterbi algorithm (Forney, 1973), and the likelihood under the null model. The score of a BGC-NRP pair is defined as the maximum score over all choices of an HMM for the BGC and a linearization for the NRP.

### 2.5. Parameter estimation

Model parameters were estimated from 145 NRP-related BGCs from MIBiG v3.1 (Terlouw *et al*., 2023) with detailed biosynthetic pathway annotations. From these entries, we manually curated 234 ground-truth BGC–NRP alignments, yielding 1,835 validated module–monomer correspondences.

These correspondences were used to calibrate PARAS probabilities by estimating how often a substrate predicted with a probability *p* was actually incorporated.

The same dataset was used to estimate transition probabilities for module skipping and insertion events, as well as conditional probabilities of methylation and L/D configuration given the presence or absence of the methylation and epimerization domains. All parameters were computed as empirical frequencies over the curated alignments.

All parameters were computed as empirical frequencies in the curated alignments.

### 2.6. Output and reporting

For each BGC–NRP pair, Nerpa 2 reports the log-odds score and the corresponding module–monomer alignment inferred from the most probable HMM state path. For each BGC and/or NRP, the top-*k* highest-scoring matches are retained, with *k* configurable by the user.

Results are provided in machine-readable JSON format to support integration into genome-mining pipelines. Additionally, an interactive HTML report summarizes top-scoring matches, including alignment details, annotated BGC domain organization, and graphical representations of the matched NRP structures and monomer graphs.

## 3. Results

### 3.1. Accuracy of BGC–NRP linking

We constructed a benchmark from MIBiG v4.0 (Zdouc *et al*., 2025), a curated database of experimentally characterized BGC– product pairs. The initial set comprised NRP-producing BGCs with at least three A domains and associated NRP structures containing at least three recognized monomers. These candidates and the 145 training BGCs from MIBiG v3.1 were clustered together using BiG-SCAPE v1.1.9 (Navarro-Muñoz *et al*., 2020). Candidate BGCs belonging to clusters containing any training BGC were excluded. This filtering yielded 200 BGCs and 367 associated NRPs. The structural database was further extended with Norine entries and deduplicated, yielding 1,058 non-isomorphic NRP monomer graphs.

We compared Nerpa 2 (v2.1.0) against Nerpa 1 (Kunyavskaya *et al*., 2021) (v1.1.0, updated for compatibility with antiSMASH v7 (Blin *et al*., 2023)) and BioCAT v1.0.0 (Konanov *et al*., 2022).

Each BGC was scored against all candidate NRPs (211,600 BGC–NRP comparisons), retaining the 10 highest-scoring predictions per BGC for each tool. A BGC was considered correctly identified at rank *k* if at least one ground-truth product of the BGC appeared within the top *k* NRP predictions.

As shown in Fig. 3a, Nerpa 2 consistently outperforms Nerpa 1 and BioCAT across ranking thresholds. At rank 1, Nerpa 2 recovered 47.5% of annotated products, compared to 39.5% for Nerpa 1 and 15.0% for BioCAT. At rank 10, recovery increased to 77.5%, exceeding 59.0% and 35.5% for Nerpa 1 and BioCAT, respectively.

**Figure 3:**
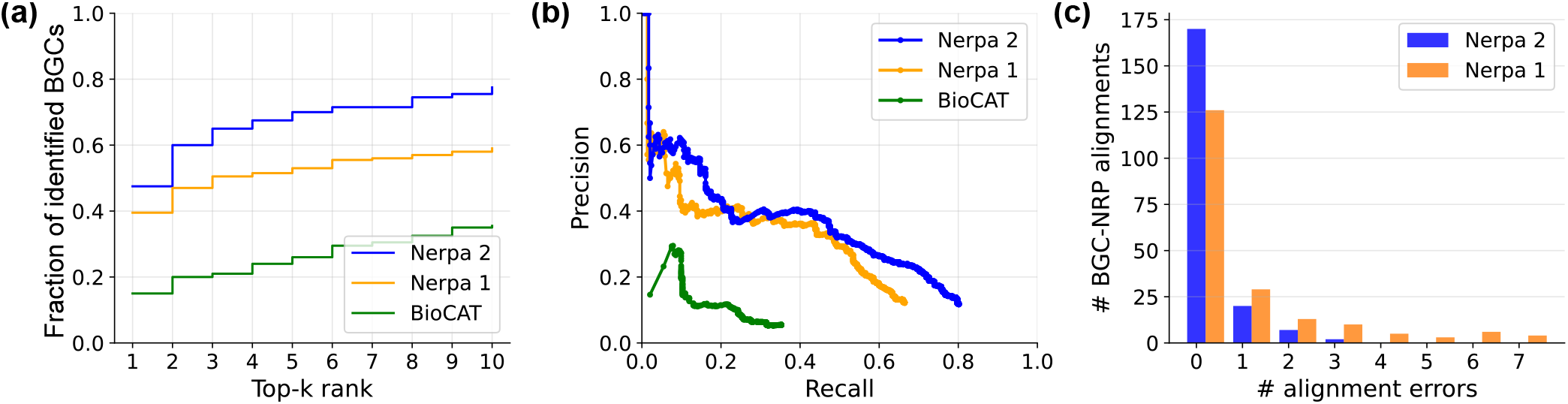
BGC–NRP linking and alignment performance on MIBiG-derived benchmarks. (a) Rank-based recovery of ground-truth products: fraction of BGCs with at least one annotated product within the top *k* ranked NRP predictions. (b) Precision–recall analysis over the top 10 predictions per BGC, treating annotated BGC–NRP pairs as positives and all other pairs as negatives. (c) Distribution of module–monomer alignment errors on ground-truth BGC–NRP alignments.

We further evaluated how well the scoring functions distinguish correct from incorrect BGC–NRP links using precision–recall analysis over the top 10 predictions per BGC (Fig. 3b). The ground-truth BGC–NRP pairs from MIBiG were treated as positives and all other pairs as negatives. Nerpa 2 achieves higher precision than Nerpa 1 across most of the recall range, indicating a more favorable trade-off between recovering correct links and introducing false positives. This advantage is most pronounced at moderate recall (*>* 0.3), while Nerpa 1 shows a slight edge at very low recall (*<* 0.05), corresponding to the highest-confidence predictions. BioCAT exhibits substantially lower precision across the entire range.

### 3.2. Correctness of module–monomer alignments

Unlike BioCAT, Nerpa produces explicit module–monomer alignments, enabling direct assessment of biosynthetic pathway reconstruction. We evaluated this using the manually curated dataset comprising 234 ground-truth BGC–NRP alignments (Section 2.5). Because this dataset was also used to train Nerpa 2, we retrained the model for this experiment using five-fold cross-validation. To reduce similarity between folds, BGCs were partitioned based on BiG-SCAPE v1.1.9 similarity. Nerpa 1 was applied without retraining, providing it with a favorable evaluation setting due to partial overlap with its training data (Kunyavskaya *et al*., 2021).

An alignment error was defined as any deviation from the curated module–monomer correspondence, including incorrect monomer assignment, failure to skip a module, or erroneous stuttering or insertion events. As shown in Fig. 3c, Nerpa 2 yields substantially more perfectly reconstructed alignments than Nerpa 1 (170 vs 126). The total number of alignment errors decreases from 184 to 40, reflecting improved handling of non-collinear and irregular assembly lines.

Representative examples of an iterative and a module-skipping assembly line are shown in Supplementary Fig. S1 and S2. Both tools identified the correct NRP, but Nerpa 1 failed to fully reconstruct module–monomer correspondences, whereas Nerpa 2 produced error-free alignments.

### 3.3. Scalability and practical applicability

We screened all genomes in the antiSMASH database v5 (Blin *et al*., 2026) containing predicted NRP-related BGCs with at least three A domains (17,305 genomes, 116,054 BGCs) against pNRPdb v2, an updated database of known and putative NRP structures compiled in this study, comprising 4,972 non-isomorphic monomer graphs. This resulted in over 5 *×* 10^8^ BGC–NRP comparisons. The full analysis completed in 9 hours using 50 CPU threads on an AMD Opteron 6378 2.4 GHz compute node. The top 10 predictions per BGC were retained, and the 10,000 highest-scoring BGC–NRP matches overall were selected for downstream analysis.

Because BGCs in the antiSMASH database lack curated product annotations, we assessed prediction plausibility using genus-level consistency. Many NRPs are taxonomically restricted; therefore, agreement between the genus of the BGC-containing genome and the reported producer genus of the matched compound provides supporting biological context. Importantly, Nerpa 2 does not incorporate taxonomic information or global sequence similarity into its scoring model, making genus-level agreement an orthogonal line of evidence. In this large-scale screening, genus agreement reached 84% among the top 100 ranked BGC–NRP pairs and decreased to 40% at rank 10,000, yet remained consistently above the random matching baseline of 7.5% (Supplementary Fig. S3).

We further examined the 20 highest-scoring BGC–NRP pairs under the constraint of one best NRP per BGC and one best BGC per NRP. For each pair, we identified an appropriate reference BGC corresponding to the Nerpa-matched compound: in seven cases, this was the exact MIBiG BGC, in eleven cases, a related MIBiG BGC producing a structurally similar compound, and in two cases (ramoplanin A1 and paenialvin A), a literature-derived BGC not present in MIBiG (Supplementary Table S1).

All 20 BGC–NRP pairs were evaluated using cblaster (Gilchrist *et al*., 2021), a sequence-based BGC comparison tool, against MIBiG v4.0 supplemented with the two missing reference BGCs. In the vast majority of cases, the reference BGC corresponding to the Nerpa-matched compound ranked first or exhibited near-identical similarity to the top cblaster hit (Supplementary Table S1), providing orthogonal sequence-based support for the structure-based assignments.

For ramoplanin A1 and paenialvin A, the Nerpa-predicted BGCs demonstrated substantially higher similarity than any MIBiG entry, suggesting that these represent plausible gene clusters not yet curated in MIBiG (Supplementary Fig. S4). Notably, the original publications described only the compound structures and producing species (Ciabatti *et al*., 1989; Meng *et al*., 2018); the ramoplanin BGC was reported decades later (Yushchuk *et al*., 2024), while the paenialvin BGC remains uncharacterized.

## 4. Discussion

Linking biosynthetic gene clusters (BGCs) to their chemical products remains a central challenge in genome mining. Here, we presented a probabilistic framework addressing this problem for nonribosomal peptides (NRPs). By explicitly modeling substrate selection uncertainty and diverse biosynthetic pathways within a unified HMM formulation, Nerpa 2 improves robustness in complex and non-collinear NRP assembly lines while remaining scalable to large genome and compound databases.

On curated benchmarks, Nerpa 2 improves the accuracy of BGC–NRP linking and more reliably reconstructs the underlying biosynthetic logic. Importantly, the tool does not rely on sequence similarity or taxonomy, enabling complementary validation of the reported BGC annotations.

Beyond benchmarking performance, our approach supports large-scale genome mining and community curation efforts such as MIBiG “Annotathons”. In our antiSMASH database screening, structure-based linking highlighted plausible candidate BGCs for compounds lacking curated annotations, including recently isolated peptide antibiotic paenialvin A. Nerpa 2 facilitates dereplication of BGCs likely associated with known compounds and prioritization of putatively novel BGCs. More broadly, the interpretable links between BGCs and product structures provide a foundation for advancing our understanding of NRP biosynthesis and informing combinatorial bioengineering of novel compounds.

## Acknowledgments

We thank Dr. Kenan Bozhüyük for insightful discussions on NRP biosynthesis and bioengineering applications of Nerpa 2, and Petr Popov for exploring alternative scoring and speed-up strategies.

## Supplementary data

Supplementary data are available online.

## Data availability

Nerpa (v2.1.0 and v1.1.0), pNRPdb v2, curated training data, materials and scripts required to reproduce the results are freely available at https://github.com/gurevichlab/nerpa. MIBiG BGC–NRP pairs, Norine structures, and antiSMASH database BGCs were obtained from their respective public repositories. Detailed results of the antiSMASH database v5 screening, including the top-20 manually analyzed cases and a minimal report of the top-10,000 BGC–NRP matches, are available at Zenodo (DOI: 10.5281/zenodo.19048303).

