## Supplementary data for "Nerpa 2: probabilistic linking of biosynthetic gene clusters to nonribosomal peptides"

### Supplementary Material for “Nerpa 2: probabilistic linking of biosynthetic gene clusters to nonribosomal peptides”

### Supplementary Note 1: processing of A domain specificity predictions

This section describes how Nerpa 2 adjusts the probability distributions reported by PARAS [1]. The goal is to make the probabilities reflect the actual chance that a prediction is correct. This makes the probabilities easier to interpret and tends to improve downstream inference.

Define:

- $A_{\text{PARAS}}$ — the set of PARAS-supported amino acids.
- $A_{Nerpa2}$ — the set of Nerpa 2-supported amino acids (core residues).
- $\texttt{CORE\_RES} : A_{\text{PARAS}} \to A_{Nerpa2}$ — a surjective conversion function.
- $P_{\text{PARAS}} : A_{\text{PARAS}} \to [0,1]$ — the probability distribution reported by PARAS.
- $C : [0,1] \to [0,1]$ — the calibration function learned during training.
- $P_{\text{null}} : A_{Nerpa2} \to [0,1]$ — the null-hypothesis distribution.
- $\varepsilon \in [0,1]$ — the null-mixing weight (0.05 by default).

Then the adjusted probability distribution $P_{Nerpa2}$ is obtained as follows:

1. **Convert PARAS amino acids to Nerpa 2 core residues (merge probability mass).**

$$\forall r \in A_{Nerpa2} : \quad P_{\text{raw}}(r) \leftarrow \sum_{\substack{a \in A_{\text{PARAS}} \\ \texttt{CORE\_RES}(a)=r}} P_{\text{PARAS}}(a)$$

2. **Calibrate.**

$$\forall r \in A_{Nerpa2} : \quad P_{\text{cal}}(r) \leftarrow C(P_{\text{raw}}(r))$$

3. **Normalize (mix with null distribution total confidence is below 1).**

$$Total \leftarrow \sum_{r \in A_{Nerpa2}} P_{\text{cal}}(r)$$

$$\forall r \in A_{Nerpa2} : \quad P_{norm}(r) \leftarrow \begin{cases} \dfrac{P_{\text{cal}}(r)}{Total} & \text{if } Total > 1, \\ P_{\text{cal}}(r) + (1 - Total) \cdot P_{\text{null}}(r) & \text{otherwise.} \end{cases}$$

4. **Mix with a small portion of the null distribution to avoid zero probabilities**.

$$\forall r \in A_{Nerpa2}: \quad P_{\text{Nerpa 2}}(r) \;\leftarrow\; (1-\varepsilon)\cdot P_{norm}(r) + \varepsilon \cdot P_{\text{null}}(r)$$

### Supplementary Note 2: generation of tentative assembly lines

We generate candidate assembly-line orders based on genomic coordinates, strand information, and domain composition, allowing some contiguous blocks of genes to be flipped when domain-architecture heuristics implies a possibility of non-collinearity.

1. **Form strand blocks.** We group adjacent core genes on the same DNA strand into *blocks*. Blocks are kept contiguous when constructing candidate assembly-line orders.

2. **Fix or flip within-block order using domain heuristics.** For each block, we test whether its within-block gene order is *consistent* with simple domain-knowledge constraints on module position (e.g., a starter condensation domain should occur only in the first module, and a termination thioesterase domain only in the last module). If a block is consistent, we keep its within-block order as annotated. If it is inconsistent, we also test the reversed within-block order (i.e., ordering genes in the opposite transcription direction). If the reversal is consistent, we mark the block as *flippable* and allow both within-block orientations in downstream candidate generation.

3. **Generate tentative gene orders.** Using the resulting within-block orders (fixed blocks) and within-block options (flippable blocks), we concatenate blocks in (i) increasing genomic coordinate and (ii) decreasing genomic coordinate. For each of these two concatenations, we enumerate all choices of orientations for flippable blocks. If there are $k$ flippable blocks, this produces $2 \cdot 2^k$ tentative gene orders.

4. **Extract A-domain-bearing modules.** For each tentative gene order, we extract all modules containing A domains and list them in order. Each module records its immediate genomic context, such as its domain composition,

presence of PKS domains upstream/downstream and its position within the block.

Note that the resulting tentative assembly lines incorporate all core genes containing an A domain and identified by antiSMASH, some of which could be artifacts or belong to neighboring BGCs. To account for this, we retain the information about the borders of gene blocks and have small penalties for skipping entire gene blocks at the ends of a tentative assembly line in subsequent alignment against NRP linearizations.

This approach allows us to handle complex assembly lines effectively while minimizing the impact of unrelated or misclassified core genes in the BGC.

**Supplementary Table S1:** Top 20 BGC–NRP matches identified by Nerpa 2 in the screening of the antiSMASH database v5 against pNRPdb v2. For each match, a reference BGC corresponding to the matched NRP was identified and the pair was further evaluated using `cblaster` [2] against MIBiG v4.0 supplemented with the missing reference BGCs. The tool was run with default parameters and a minimum score cutoff of 10.0. The columns $S$(Ref) and $S$(MIBiG) show `cblaster` scores of the identified BGCs against the corresponding reference BGC and against the best-scoring BGC in MIBiG, respectively; "n/a" indicates that no hit was detected. If the best MIBiG hit differs from the reference BGC, the higher of the two scores is shown in bold. The complete Nerpa HTML report for these matches, GenBank files for 20 identified BGCs and the two reference BGCs missing in MIBiG, are available at Zenodo `https://doi.org/10.5281/zenodo.19048303`. Genus abbreviations: *A. Actinoplanes*; *M. Micromonospora*; *P. Pseudomonas*; *S. Streptomyces*; *B. Bacillus*; *Bb. Brevibacillus*; *Pb. Paenibacillus*; *Pr. Photorhabdus*; *Xr. Xenorhabdus.*

| Nerpa Match | | Matched BGC | | Matched NRP | | | `cblaster` | | |
|---|---|---|---|---|---|---|---|---|---|
| ID | Score | Genome ID | Organism | NRP ID | Name | Producer | Reference BGC | $S$(Ref) | $S$(MIBiG) |
| 000 | 53.72 | GCF_013385425.1 | *P. gingeri* | BGC0001629-0 | jessenipeptin | *P.* sp. QS1027 | BGC0001629 (jessenipeptin) | n/a[a] | **35.70** |
| 001 | 51.22 | GCF_001705995.2 | *P.* sp. BIOMIG1BAC | BGC0000425-0 | sessilin A | *P.* sp. CMR12a | BGC0000425 (sessilin A) | 21.41 | 21.41 |
| 002 | 50.68 | GCF_044333315.1 | *S. exfoliatus* | NPA007111 | CB-182350 | *S. fradiae* | BGC0000291 (A54145)[i] | 29.38 | 29.38 |
| 003 | 50.58 | GCF_036213785.1 | *Bb. parabrevis* | BGC0002832-0 | C15-gramicidin | *Bb. parabrevis* HD1.4A | BGC0000367 (gramicidin)[ii] | 23.61 | 23.61 |
| 004 | 48.82 | GCF_030687655.1 | *P. tolaasii* | NPA020327 | tolaasin I | *P. tolaasii* | BGC0000447 (tolaasin I/F) | 17.04[b] | **22.66**[b] |
| 005 | 48.79 | GCF_022836895.1 | *P. palleroniana* | BGC0000447-1 | tolaasin F | *P. costantinii* | BGC0000447 (tolaasin I/F) | 18.51[b] | **20.67**[b] |
| 006 | 48.23 | GCF_002865525.1 | *Bb. laterosporus* | BGC0002432-0 | laterocidine | *Bb. laterosporus* LMG 15441 | BGC0002432 (laterocidine) | 64.93 | 64.93 |
| 007 | 48.13 | GCF_038593055.1 | *Pb.* sp. FSL L8-0502 | NPA015860 | tridecaptin A1 | *Pb. terrae* NRRL B-30644 | BGC0000449 (tridecaptin) | 12.42[c] | 12.42[c] |
| 008 | 47.46 | GCF_044327715.1 | *S. fungicidicus* | NOR01908 | enduracidin A' | *S. fungicidicus* | BGC0000341 (enduracidin A/B) | 53.93 | 53.93 |
| 009 | 47.29 | GCF_007831095.1 | *B. paralicheniformis* | BGC0000310-0 | bacitracin A1 | *B. licheniformis* | BGC0000310 (bacitracin) | 19.98 | 19.98 |
| 010 | 46.49 | GCF_000196155.1 | *Pr. laumondii* TTO1 | NPA020256 | kolossin A | *Pr. luminescens* | BGC0001641 (kolossin) | n/a[d] | n/a[d] |
| 011 | 46.46 | GCF_002706795.1 | *Bb. laterosporus* DSM 25 | NPA028417 | brevicidine | *B. laterosporus* DSM 25 | BGC0001536 (brevicidine) | 24.78[e] | **47.91**[e] |
| 012 | 45.92 | GCF_043198755.1 | *M. chersina* NPDC048726 | NPA028646 | ramoplanin A1 | *A.* sp. ATCC 33076 | PP681430.1.region001[iii] | **46.19** | 30.72 |
| 013 | 45.62 | GCF_003171815.2 | *B. paralicheniformis* | NOR00020 | bacitracin B2 | *B. licheniformis* | BGC0000310 (bacitracin) | 19.97 | 19.97 |
| 014 | 45.18 | GCF_004794665.1 | *S.* sp. S816 | NPA023003 | actinomycin D1 | *S.* sp. LHW52447 | BGC0000296 (actinomycin D) | 48.04 | 48.04 |
| 015 | 44.22 | GCF_900155355.1 | *Xr. innexi* | BGC0000464-1 | xenoamicin B | *Xr. doucetiae* | BGC0000464 (xenoamicin A/B) | 18.65 | 18.65 |
| 016 | 44.06 | GCF_022832475.1 | *B.* sp. B19-2 | NPA012256 | bacitracin B2 | *B.* sp. SNA-60-367 | BGC0000310 (bacitracin) | 19.98 | 19.98 |
| 017 | 44.05 | GCF_044330875.1 | *S. roseolus* | NPA020427 | A-54145 F | *S. fradiae* | BGC0000291 (A54145) | 30.45 | 30.45 |
| 018 | 43.61 | GCF_025547055.2 | *Pb. alvei* | NPA021780 | paenialvin A | *Pb. alvei* DSM 29 | AMBZ01000007.1.region002[iv] | **81.66** | n/a |
| 019 | 43.20 | GCF_000156695.2 | *S. filamentosus* NRRL 11379 | BGC0000336-0 | daptomycin | *S. filamentosus* NRRL 11379 | BGC0000336 (daptomycin) | 33.88 | 33.88 |

Notes on the reference BGC selection:

[i] For NPA007111 (CB-182350), BGC0000291 (A54145) was used as the reference because these NRPs belong to the same compound family [3].

[ii] For BGC0002832-0 (C15-gramicidin), BGC0000367 (gramicidin) was used because BGC0002832 was retired and the corresponding BGC sequence is no longer available.

[iii] For NPA028646 (ramoplanin A1), the reference BGC was taken from [4] as no related entry is currently available in MIBiG.

[iv] For NPA021780 (paenialvin A), no experimentally characterized BGC is available; the genome of the producing organism *Paenibacillus alvei* DSM 29 was analyzed with antiSMASH v8 [5], and the only NRP-related BGC with the matching number of A domains was selected as a putative reference.

Notes on low and missing `cblaster` scores:

[a] BGC0001629 (jessenipeptin) contains only two genes and therefore cannot yield a meaningful `cblaster` score by design.

[b] The best MIBiG hit is BGC0002768 (asplenin), which lacks an associated compound structure and is therefore absent from pNRPdb and the Nerpa search space.

[c] BGC0000449 (tridecaptin) contains only five genes, leading to a relatively low score despite 90% sequence identity between the NRP-encoding genes.

[d] BGC0001641 (kolossin) contains only a single gene and therefore cannot produce a meaningful `cblaster` score.

[e] BGC0001536 (brevicidine) is shorter (11 genes) than the best-scoring hit BGC0002432 (laterocidine, 36 genes), resulting in a lower score despite 100% identity of the NRP-encoding genes, while the corresponding genes in the longer hit show only 57%–88% identity.

### Supplementary Figures

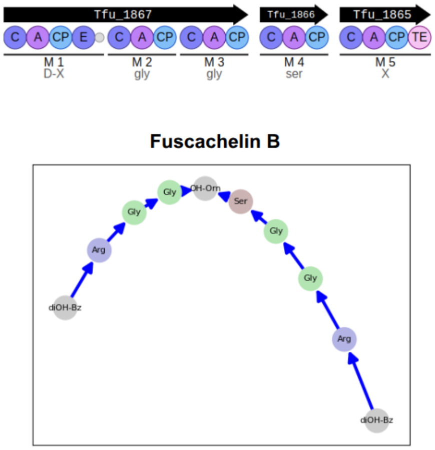


**Nerpa 1**

| BGC Module | Monomer |
|---|---|
| --- | diOH-Bz |
| Tfu_1867_1 | Arg |
| Tfu_1867_2 | Gly |
| Tfu_1867_3 | Gly |
| Tfu_1866_1 | Ser |
| Tfu_1865_1 | --- |

**Nerpa 2**

| BGC Module | Monomer | Score |
|---|---|---|
| --- | diOH-Bz | -2.5 |
| Tfu_1867_1 | Arg | 3.754 |
| Tfu_1867_2 | Gly | 2.325 |
| Tfu_1867_3 | Gly | 2.324 |
| Tfu_1866_1 | Ser | 2.445 |
| Tfu_1865_1 | OH-Orn | 3.373 |
| --- | diOH-Bz | -2.5 |
| Tfu_1867_1 | Arg | 3.754 |
| Tfu_1867_2 | Gly | 2.325 |
| Tfu_1867_3 | Gly | 2.324 |
| Tfu_1866_1 | --- | -2.5 |
| Tfu_1865_1 | --- | -2.5 |

**Supplementary Figure S1:** Alignments produced by Nerpa 1 and Nerpa 2 for an iterative BGC that produces Fuscachelin B (MIBiG:BGC0000359; ground-truth alignment derived from [6]). This BGC produces two similar fragments that are subsequently merged. Nerpa 2 successfully identified the iterative nature of the BGC and linked all NRP monomers to the corresponding BGC modules, while Nerpa 1 was able to match only half of the NRP monomers to the respective modules.

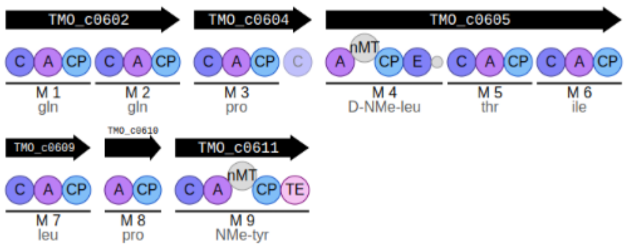


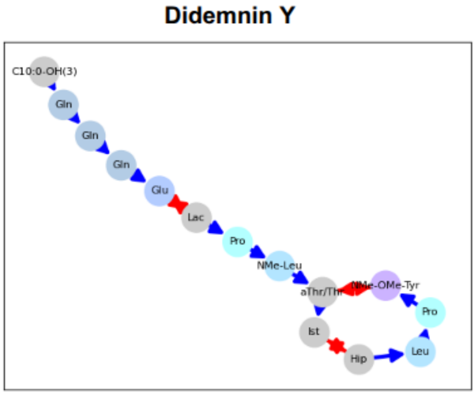


**Nerpa 1**

| BGC Module | Monomer |
|---|---|
| TMO_c0602_1 | Gln |
| TMO_c0602_2 | Gln |
| --- | Gln |
| --- | Glu |
| --- | Lac |
| TMO_c0604_1 | Pro |
| TMO_c0605_1 | NMe-Leu |
| TMO_c0605_2 | aThr |
| --- | Ist |
| TMO_c0605_3 | Hip |
| TMO_c0609_1 | Leu |
| --- | Pro |
| --- | NMe-OMe-Tyr |

**Nerpa 2**

| BGC Module | Monomer | Score |
|---|---|---|
| TMO_c0602_1 | Gln | 2.492 |
| TMO_c0602_2 | Gln | 2.491 |
| TMO_c0602_1 | Gln | 2.492 |
| TMO_c0602_2 | Glu | -0.789 |
| --- | Lac | -2.5 |
| TMO_c0604_1 | Pro | 1.994 |
| TMO_c0605_1 | NMe-Leu | 1.609 |
| TMO_c0605_2 | aThr/Thr | 2.796 |
| TMO_c0605_3 | Ist | -1.797 |
| --- | Hip | -2 |
| TMO_c0609_1 | Leu | 1.589 |
| TMO_c0610_1 | Pro | 1.994 |
| TMO_c0611_1 | NMe-OMe-Tyr | 2.831 |

**Supplementary Figure S2:** Alignments produced by Nerpa 1 and Nerpa 2 for the BGC that produces Didemnin Y (MIBiG:BGC0000985; ground-truth alignment derived from [7]), which has a non-standard assembly line. The first gene is iterative, and the BGC includes unusual features, such as several modules lacking condensation (C) domains, which are typically only absent at the beginning of the assembly line. Nerpa 2 correctly captured the iteration of the first gene and constructed the entire assembly line, while Nerpa 1 failed to do so. Nerpa 1 missed the iteration and was unable to handle multiple modules without C domains, resulting in an incomplete alignment.

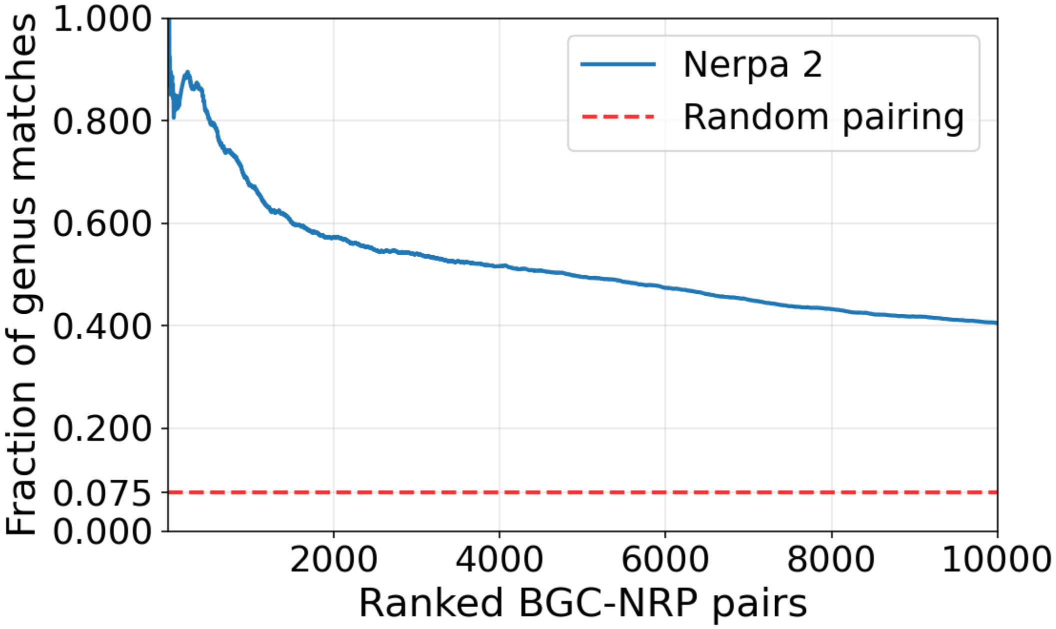


**Supplementary Figure S3:** Cumulative genus-level agreement among ranked BGC–NRP pairs. Cumulative fraction of pairs where the BGC genome genus matches the compound's reported producer genus, plotted versus rank. The dashed red line is the random-pairing baseline, $\sum_g p_g q_g$, where $p_g$ and $q_g$ are genus frequencies among BGCs and NRPs, respectively.

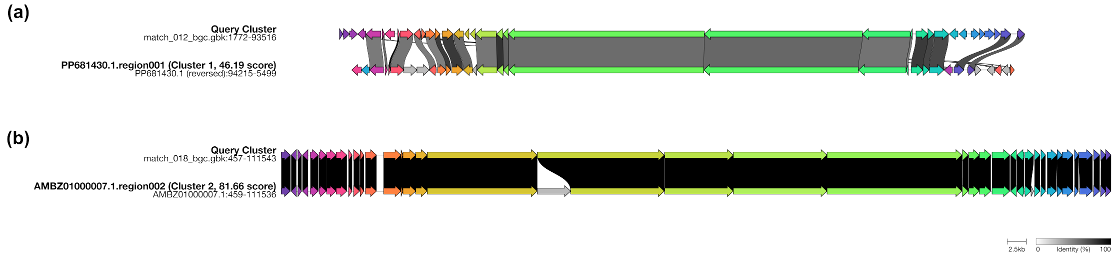


**Supplementary Figure S4:** `clinker` [2] visualization of homology between Nerpa-identified candidate BGCs and reference BGCs. **a.** Candidate ramoplanin A1 BGC identified in *M. chersina* (GCF_043198755.1) compared to the reference ramoplanin BGC characterized in *Actinoplanes* sp. ATCC 33076 [4]. **b.** Candidate paenialvin A BGC identified in *P. alvei* (GCF_025547055.2) compared to the putative reference cluster identified in the genome of the paenialvin-producing strain *Paenibacillus alvei* DSM 29.
